# Strong genotype-by-genotype and genotype-by-environment interactions induced by antibiotic resistance mutations

**DOI:** 10.64898/2025.12.03.692229

**Authors:** Alicia H. Williams, Ian Gooi, Madelaine Boggan, Ziqing Liu, Andrew D. Letten, Jan Engelstädter

## Abstract

The often parallel evolution of antibiotic resistance in different species offers a promising basis for predicting evolutionary change. However, the extent to which parallel genetic evolution is reflected at the phenotypic level remains unclear. We investigated how genetic background and temperature influence the fitness effects of homologous rifampicin resistance mutations in absence of antibiotics across two temperatures and three species (*Escherichia coli*, *Acinetobacter baylyi* and *Lactiplantibacillus plantarum*). We measured the effect of resistance mutations on two growth traits (maximum growth rate and maximum optical density) across all conditions and fit linear models to this data using Bayesian inference. Our model fits provide evidence for pervasive genetic interactions in both growth traits, including mutation by species (GxG), mutation by temperature (GxE), and GxGxE interactions. In particular, the effect of resistance mutations was strongly dependent on genomic background, to the extent that mutations often reduced growth in one species but increased it in another (sign epistasis). Similarly, the effect of mutations on growth was strongly influenced by temperature, with some mutations increasing growth at one but decreasing growth at the other temperature. Most notably, three resistance mutations induced strong heat sensitivity in *A. baylyi*. Our results reinforce previous studies that report strong epistatic and GxE effects of resistance mutations, indicating that in the absence of additional data and more mechanistic models, predicting fitness effects of resistance mutations across species and environments can be very challenging.

## Introduction

Owing to theoretical advances and technological innovation, recent years have seen a surge in interest in predicting evolutionary change (Lassig et al. 2017; Wortel et al. 2023). Accurate predictions could benefit fields in which evolution is rapid and its impacts significant, including medicine, public health, agriculture, and biological conservation. In particular, forecasting antibiotic resistance evolution in bacteria has received considerable attention (Schenk and de Visser 2013; Pinheiro 2024; Rolff et al. 2024). This is not only because of the stakes involved given the emerging resistance crisis, but also because resistance mutations hold promise for successful predictions, as antibiotic resistance often has a relatively simple genetic basis that is highly conserved across species (Wong and Kassen 2011). This is especially true for mutations at the antibiotic target site that confer resistance by preventing antibiotic binding (Blair et al. 2015). Since antibiotic targets are usually proteins involved in core cellular functions common to all bacteria (e.g., protein synthesis), resistance often evolves via homologous mutations in different species. This parallelism offers the promise of predictions that are robust with regard to species or strain.

Resistance to rifampicin (RIF) is one such case. RIF is an antibiotic that has been used since the 1960s to treat a wide range of infections and is a first-line drug to treat tuberculosis (Goldstein 2014; Adams et al. 2021). RIF binds to the *β*-subunit of the RNA polymerase (RNAP), blocking elongation of newly formed transcripts. Resistance to rifampicin typically evolves through point mutations in the gene *rpoB* that codes for the RNAP *β*-subunit. Early mutant screens in *E. coli* demonstrated that these mutations typically arise at specific positions in *rpoB* that lead to amino acid changes close to the binding site of RIF (Jin and Gross 1988), and subsequent work has demonstrated that the same (i.e., homologous) mutations also confer RIF resistance in *Mycobacterium tuberculosis* and many other species (e.g. Morlock et al. 2000; Goldstein 2014). Given the broad phylogenetic distribution of these species, RIF resistance evolution presents a remarkable case of parallel evolution across the entire bacterial tree of life (Bolourchi et al. 2025).

It is less clear to what extent parallel evolution of antibiotic resistance at the genetic level is reflected at the phenotypic level. Antibiotic resistance is typically associated with fitness effects in the absence of drugs, measured for example as a change in the growth rate or competitive fitness relative to the ancestral genotype. (This is often referred to as the ‘cost of resistance’, but here we follow Lenormand et al. (2018) and use the more neutral term ‘fitness effects in absence of drugs’.) Our ability to predict the evolution of antibiotic resistance depends on these fitness effects being similar across different genetic backgrounds (different strains or different species), and also across different environments. However, there is mounting evidence that fitness effects can vary significantly by both genetic background (Vogwill et al. 2016; Wong 2017; Vogwill and MacLean 2015; Vogwill et al. 2014; Trindade et al. 2009; Apjok et al. 2019; Hinz et al. 2024; Rodŕıguez-Verdugo et al. 2013; Borrell et al. 2013) and environment (Clarke et al. 2020; Leehan and Nicholson 2021, 2022; Gifford et al. 2016; Mira et al. 2022; Ghenu et al. 2023). Variation of fitness effects by genetic background can be considered a case of epistasis, also referred to as genotype-by-genotype (G×G) interactions. Epistasis is defined as a deviation from independent genetic effects (here, effects of resistance mutations and species/strain on fitness). Independence can be defined on either an additive or multiplicative scale. In extreme cases, deviations from independence can be so pronounced that not only the magnitude of a phenotypic effect depends on the genetic background (magnitude epistasis) but also its direction (sign epistasis; Weinreich et al. 2005). Variation of fitness effects in different environments are referred to as a genotype-by-environment (G×E) interactions, considered by some authors as a special case of pleiotropy (but see Lenormand et al. 2018). Conceptually, GxE effects are analogous to GxG effects and we can again distinguish between magnitude and sign GxE effects (or pleiotropy). A final complication occurs when epistasis itself is dependent on the environment (or, equivalently, GxE effects depend on genetic background), giving rise to GxGxE interactions.

Here we investigated the impact of genetic background and temperature on the fitness effects of RIF resistant mutants in the absence of antibiotics. We measured growth effects of rifampicin resistance mutations across three species, *Escherichia coli*, *Acinetobacter baylyi* and *Lactiplan-tibacillus plantarum*, and at two temperatures, 30 *^◦^*C and 37 *^◦^*C. We then fit a series of Bayesian linear models to our data, demonstrating strong GxG (epistasis), GxE, and GxGxE effects in this system.

## Methods

### Strains and culture methods

The species used were *Escherichia coli* MG1655, *Acinetobacter baylyi* ADP1 and *Lactiplan-tibacillus plantarum*. Aside from these ancestral strains (henceforth referred to as wildtype), we used a total of 55 rifampicin resistant mutants across these three species that were obtained in prior mutant screens (Gooi et al., in preparation). *E. coli* and *A. baylyi* were grown in Luria-Bertani (LB) broth, and *L. plantarum* (which is unable to grow in LB) was grown in De Man–Rogosa–Sharpe (MRS) broth. Unless specified, species were incubated in their recorded optimal temperature; 37*^◦^*C for *E. coli* and 30*^◦^*C for *A. baylyi* and *L. plantarum*. Stocks of all strains and mutants were stored at −80*^◦^*C in 15% glycerol within the preferred media for the species. Overnight cultures used for the following experiments were grown from either a scraping of the −80*^◦^*C glycerol stock or picked from a colony on LB agar spread from the −80*^◦^*C glycerol stock.

### Growth assays

We performed growth assays of the three wildtypes and all mutants at each species’ preferred temperature. In addition, for the *E. coli* and *A. baylyi* wildtype and all mutants with homolo-gous RIF resistance mutations across those two species we also measured growth at the other species’ preferred temperature (30 *^◦^*C and 37 *^◦^*C, respectively). For each growth assay, a 96 well plate with 178µL of growth media was inoculated with 2µL of overnight cultures. Plates were incubated under continuous shaking for 24 hours in a microplate reader (BioTec Synergy H1 or Epoch) and the absorbance optical density at 600 nm (OD_600_) was measured every 5 minutes. Growth assays were performed with six replicates for each wildtype and mutant in their preferred temperature, and with three replicates for the additional growth assays in the non-preferred temperature.

### Statistical analysis

Growth curves of all conditions from the absorbance OD_600_ measures were analysed using the in-house R package *grow96* (https://github.com/JanEngelstaedter/grow96). Two growth traits were estimated for each growth curve: maximum OD (maxOD) and the maximum per capita growth rate (*µ*_max_). We used the easylinear method (Hall et al. 2014) as implemented in the *growthrates* package (Petzold 2022) to estimate maximum growth rate.

We fit linear models to the data, with either *µ*_max_ or maxOD as the response variable, and species, temperature and mutation as the independent variables. These models included: 1) models with only species and mutation as independent variables (at each species’ preferred temperature); 2) models with only temperature and mutation as independent variables (separately for each species); and 3) full models with all three independent variables for the *E. coli* / *A. baylyi* species pair. We used STAN in conjunction with the R package *brms* (Bürkner 2017) for Bayesian model fitting, assuming improper flat priors for all model coefficients. Posteriors were then extracted from the fitted models, and the mean and 95% credible intervals (as highest density intervals) were determined from the posteriors.

## Results

### Overview

We conducted growth assays of populations grown in monoculture and absence of drugs for a total of 55 RIF resistant mutants across three species (Table 1), as well as the three susceptible wildtype genotypes. From each growth curve, we estimated the maximum per captita growth rate (*µ*_max_) and the max OD. A substantial fraction of the mutations in our panel were homologous between species, allowing us to estimate interactions between mutations and genetic background (GxG interactions, or “species-level epistasis”). For the mutations that overlapped between *E. coli* and *A. baylyi* we measured growth at two temperatures (30 *^◦^*C and 37 *^◦^*C) so as to also estimate interactions between mutations and temperature (GxE interactions).

**Table 1:** Rifampicin resistance mutants used in this study. The first two columns show the amino acid position in *E. coli* coordinates and the amino acid substitution, respectively. The remaining columns show the mutations used in this study, with numbers indicating the amino acid positions in the three focal species. Mutations shown in the same row are homologues.

| Position | Mutation | *E. coli* | *A. baylyi* | *L. plantarum* |
| --- | --- | --- | --- | --- |
| 146 | V → G | 146 | 148 |  |
| 146 | V → F | 146 |  |  |
| 509 | S → R | 509 | 516 |  |
| 511 | L → S |  | 518 |  |
| 511 | L → P | 511 |  |  |
| 512 | S → P | 512 | 519 |  |
| 512 | S → F | 512 |  |  |
| 513 | Q → P | 513 |  | 477 |
| 513 | Q → K |  |  | 477 |
| 516 | D → A |  | 523 |  |
| 516 | D → G | 516 | 523 |  |
| 516 | D → N | 516 |  | 480 |
| 516 | D → E |  |  | 480 |
| 516 | D → Y |  |  | 480 |
| 518 | N → D |  | 525 |  |
| 522 | S → F | 522 |  |  |
| 526 | H → D | 526 | 533 |  |
| 526 | H → Y | 526 | 533 | 490 |
| 526 | H → N | 526 | 533 | 490 |
| 529 | R → G | 529 |  | 493 |
| 529 | R → C | 529 |  | 493 |
| 529 | R → L | 529 |  |  |
| 529 | R → H |  |  | 493 |
| 531 | dup |  | 538 |  |
| 531 | S → L |  | 538 | 495 |
| 531 | S → F | 531 |  |  |
| 533 | L → S |  | 540 |  |
| 533 | L → W |  | 540 |  |
| 534 | G → E |  |  | 498 |
| 537 | G → D |  | 544 |  |
| 564 | P → L |  | 571 | 528 |
| 572 | I → L | 572 | 579 |  |
| 572 | I → S | 572 | 579 |  |
| 572 | I → F | 572 |  |  |
| 574 | S → L |  | 581 |  |

Figure S1 shows the *µ*_max_ and max OD estimates across the three species and two temperatures. In all three species, most resistance mutations did not have a strong effect on growth rate, although some mutations reduced growth considerably (e.g. mutations at position 529 in *E. coli*). As expected, *E. coli* grew much slower and to lower densities at 30 *^◦^*C than at 37 *^◦^*C. However, in *A. baylyi* —normally grown at 30 *^◦^*C—increasing temperature to 37 *^◦^*C had a negative effect only on max OD but not on *µ*_max_.

We used Bayesian inference to fit three types of linear models to our data, corresponding to the three types of genetic interactions: 1) models of the form Trait ∼ mutation ∗ species, where Trait can be either *µ*_max_ or maxOD, to estimate GxG effects, 2) models of the form Trait ∼ mutation ∗ temperature to estimate GxE effects, and 3) models of the form Trait ∼ mutation ∗ species ∗ temperature to estimate GxG, GxE and GxGxE effects. We visualise the fits for the former two models in plots that make it easy to see both the magnitudes and types of GxG or GxE interactions (see Figure 1 for a schematic and explanation of these plots). These types include 1) magnitude epistasis (or magnitude GxE interactions), where there are positive or negative deviation from additivity but where the direction of both effects relative to a reference genotype is the same irrespective of the state at the other locus or environment, 2) sign epistasis, where the direction of one but not the other effect differs depending on the state of the other locus or environment, and 3) reciprocal sign epistasis, where the direction of both effects differs depending on the other locus or environment.

**Figure 1:**
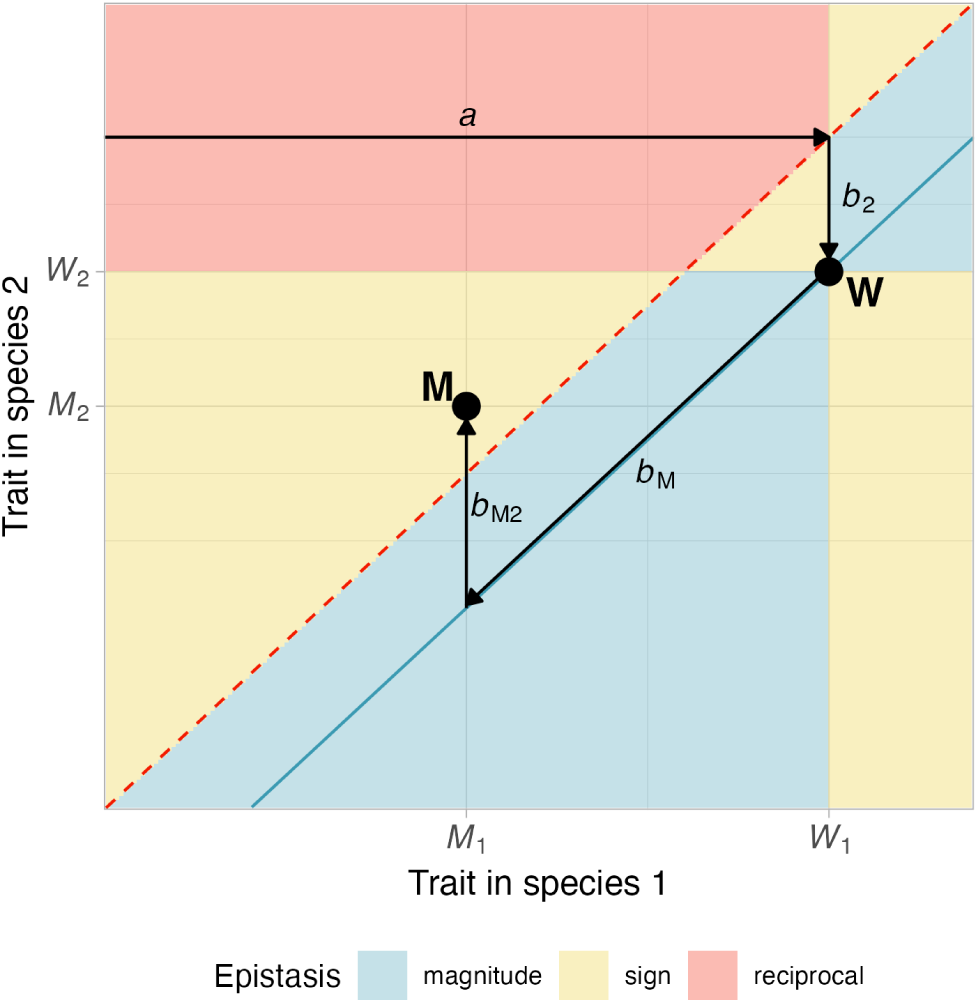
Schematic of different types of epistasis between species and resistance mutations and their relationship to linear model fits. The two axes in this plot show the values of a trait (e.g. growth rate) in species 1 and 2, respectively. The wildtype strain, carrying no resistance mutation, labelled “W”, is characterised by trait values *W*_1_ and *W*_2_ in the two species. The different areas in this plot indicate different types of epistasis (magnitude, sign and reciprocal sign epistasis) for mutants situated in these areas. The blue line indicates additive trait effects (no epistasis), and the areas above and below the blue line are characterised by positive and negative epistasis, respectively. The dashed red line indicates equal trait values of a genotype in both species. An example mutant genotype, labelled “M”, is shown that has trait values *M*_1_ and *M*_2_ in the two species. This mutant is characterised by positive (non-reciprocal) sign epistasis. This is because the mutation reduces the trait value in both species (*M*_1_ *< W*_1_ and *M*_2_ *< W*_2_) but the direction of the trait change between species 1 and 2 depends on the mutation (*W*_2_ *< W*_1_ but *M*_2_ *> M*_1_). The black arrows correspond to coefficients in a linear model fitted to data. The intercept *a* represents the baseline trait of the species 1 wildtype. Coefficient *b*_2_ captures the effect of switching from species 1 to species 2, so that *a*+*b*_2_ is the trait value for the species 2 wildtype. Coefficient *b_M_* is the effect of the mutation, so that *a* + *b*_M_ is the trait value of the species 1 mutant. Finally, *b_M_*_2_ is the coefficient capturing the interaction between species 2 and the mutation, so that *a* + *b*_2_ + *b_M_* + *b_M_*_2_ gives the trait value of the species 2 mutant. Note that “species” can be replaced by “environment” in this figure, which would then show analogous GxE instead of GxG interactions.

### Epistasis across species

Our GxG model fit reveals that epistastic interactions between RIF resistance mutations and genetic background (species) are abundant but not ubiquitous (Figure 2). For both species comparisons involving *E. coli* we see a wide spectrum of epistatic interactions affecting maximum growth rate, ranging from negative to near zero (additive effects) to positive. For example, in the *E. coli* vs. *L. plantarum* comparison (Figure 2B), mutations R529G and R529C are characterised by positive epistasis (faster growth in *L. plantarum* relative to the expectation under additive effects, given growth rates of the same mutation in *E. coli* and the two wildtypes), mutation H526Y has zero epistasis, and mutation Q513P has negative epistasis. Epistatic effects for maximum growth rate also vary considerably, although their sign appears more consistent in each species comparison. Both magnitude epistasis and, in some cases, strong sign epistasis was observed. However, sign epistasis was always unidirectional and never reciprocal. In some cases the effects of mutations at the same site (H526D vs. H526Y and R529C vs. R529G) are very similar, but this is not universally true. Overall, there does not seem to be a clear pattern for the distribution of epistatic interactions across mutations and growth traits.

**Figure 2:**
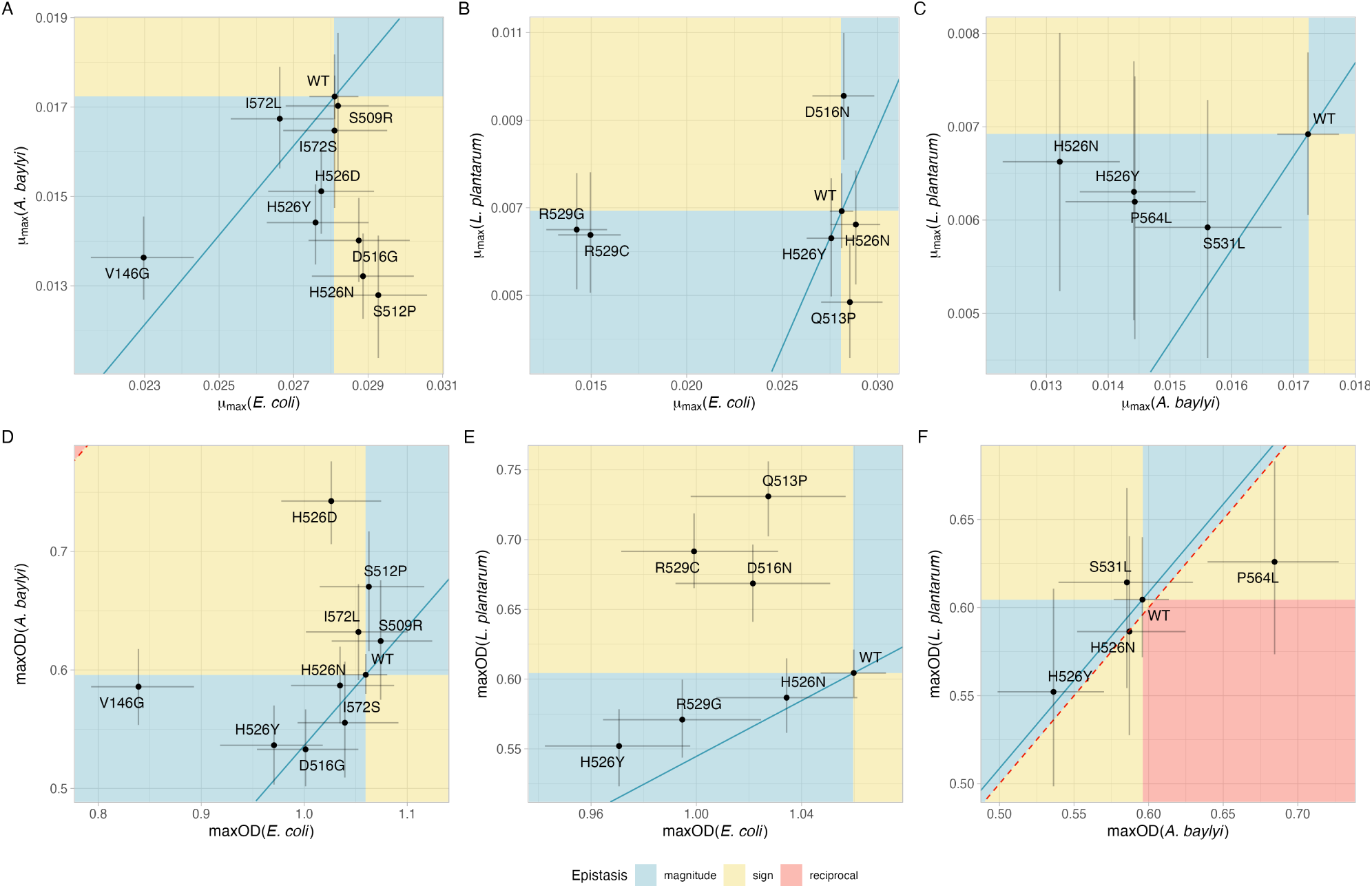
G×G interactions between rifampicin resistance mutations and species. The three columns show comparisons between three species pairs: *E. coli* vs. *A. baylyi* (A, D), *E. coli* vs. *L. plantarum* (B, E) and *A. baylyi* vs. *L. plantarum* (C, F). The two rows show the two growth traits measured: maximum growth rate (*µ*_max_, A, B, C) and maximum OD (D, E, F). In each plot, the two axes represent the estimated growth parameter for the two species. Each point shows the estimated mean of the posterior distribution of a mutant, and error bars show 95% credible intervals. The background colour denotes areas with different types of G×G interactions (magnitude, sign, or reciprocal sign epistasis). The blue lines indicates the absence of any additive epistasis (additive effects only). The dashed red line is the main diagonal and indicates equal growth trait across the two species.

### Interactions between mutations and temperature

We next investigated genetic interactions between mutations and temperature (GxE interactions), for both growth traits and the two species *E. coli* and *A. baylyi*. Figure 3A shows that in *E. coli* there are no or very weak GxE interactions involved in *µ*_max_, as seen by the overlap between the credible intervals and the blue diagonal indicating additive effects. In contrast, most mutations show strongly positive GxE effects in maxOD in *E. coli*, with many of the values falling within the domain of reciprocal sign GxE effects (red area in Figure 3C). This is because even though maxOD is considerably lower in wildtype *E. coli* at 30 *^◦^*C than at 37 *^◦^*C and is generally lower for mutants than for wildtype at 37 *^◦^*C, many mutants grow to higher maxOD than both the wildtype at 30 *^◦^*C and the same mutant at 37 *^◦^*C.

**Figure 3:**
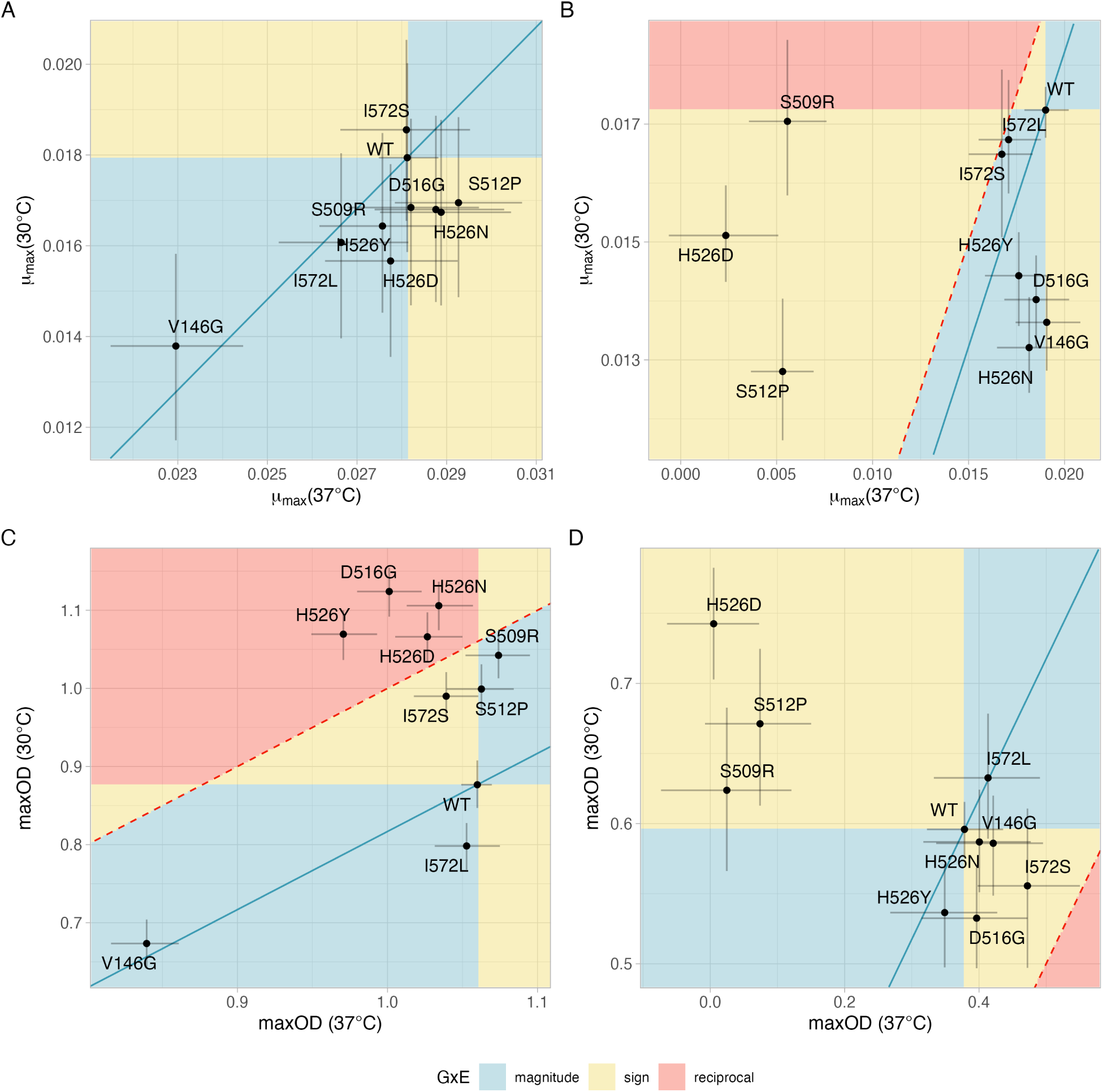
G×E interactions between rifampicin resistance mutations and temperature. The four panels correspond to the two species *E. coli* (A and C) and *A. baylyi* (B and D) as well the two growth traits maximum growth rate (*µ*_max_, A, B) and maximum OD (C, D). In each plot, the represent the estimated growth parameter at 30 *^◦^*C and 37 *^◦^*C, respectively. Each point shows the estimated mean of the posterior distribution of a mutant, and error bars show 95% credible intervals. The background colour denotes areas with different types of G×E interactions (magnitude, sign, or reciprocal sign G×E interactions). The blue lines indicates the absence of any G×E interactions (additive effects only). The dashed red line is the main diagonal and indicates growth traits across the two temperatures. See Fig. 1 for more guidance on the interpretation of this figure.

In *A. baylyi*, there is a clear pattern of three mutations (S509R, S512P and H526D) showing strongly reduced growth at 37 *^◦^*C compared to 30 *^◦^*C, both in terms of *µ*_max_ and maxOD (Figure 3B and D). These three mutations are also characterised by positive sign GxE effects because their growth is increased much more when switching from 37 *^◦^*C to 30 *^◦^*C than expected under additive effects, given the corresponding growth of the wildtype at the two temperatures. In other words, those three mutations render the bacteria particularly heat sensitive. By contrast, the other resistance mutations in *A. baylyi* exibit no or very weak GxE effects in either growth trait.

### G×G×E effects between mutations, species and temperature

We next fitted models for *µ*_max_ or max OD as a function of mutation, genetic background (species), temperature and their three-way interaction, enabling us to look at the same data as in the previous section from the angle of a full linear model. Figure 4 shows the estimated effects of the various coefficients in these model fits. These coefficients reflect estimates of epistasis (GxG), GxE effects and GxGxE effects.

**Figure 4:**
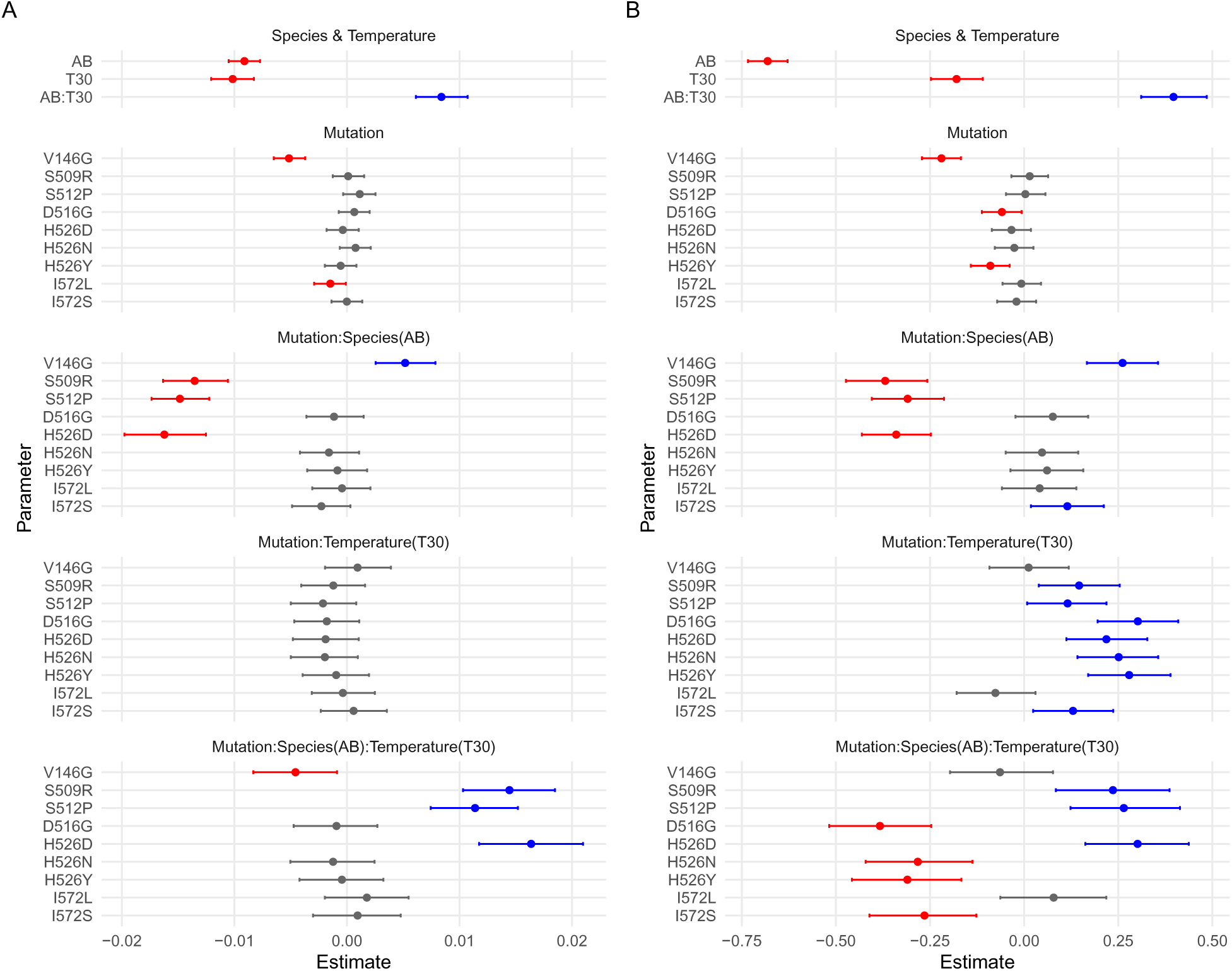
Full model fit to growth data for *E. coli* and *A. baylyi* for A) maximum growth rate and B) maximum OD. Each of the two models includes three explanatory variables (Species, Temperature and Mutation) and all possible interactions between them. The plots show, for each coefficient, the mean of the estimated posterior distribution (dots) and the 95% credible interval (horizontal lines). Coefficients with credible intervals entirely below zero are shown in red, those with credible intervals entirely above zero in blue, and those with credible intervals overlapping with zero are shown in grey. Abbreviations used for coefficients are AB=*A. baylyi* and T30=temperature of 30 *^◦^*C; colons indicate interactions terms.

The first three coefficients show the effects of species and temperature in the wildtype. The coefficients for *A. baylyi* and a temperature of 30 *^◦^*C are negative for both *µ*_max_ and maxOD, reflecting the fact that *E. coli* at 37 *^◦^*C grows better than *A. baylyi* at 37 *^◦^*C and better than at 30 *^◦^*C. However, the negative growth effects of *A. baylyi* at 37 *^◦^*C are largely compensated by the positive interaction coefficient between species and temperature (AB:T30 in Figure 4).

The effects of individual mutations, by themselves, are generally weak, with the only notable exception being V146G. This is also apparent in Figure S1 showing that most mutations (that overlapped between *E. coli* and *A. baylyi*) had only weak growth effects. Mutation by temper-ature interactions are also weak for *µ*_max_, mirroring the absence of strong GxE interactions in *E. coli* (compare Figure 3A). By contrast, there are strong mutation by species interactions for both *µ*_max_ and maxOD.

Our full model also allows us to estimate GxGxE interactions, demonstrating that such in-teractions are present for several RIF resistance mutations. These effects reflect the strong temperature sensitivity induced by the three mutations highlighted in the previous section (S509R, S512P and H526D), with the positive GxGxE coefficients for these three mutations compensating for the strongly negative coefficients describing growth at 37 *^◦^*C. To further verify the importance of GxGxE effects, we also fit a linear model that did not include the three way interaction but still included all two-way interactions (Trait ∼ mutation ∗ species + mutation ∗ temperature + species ∗ temperature instead of Trait ∼ mutation ∗ species ∗ temperature. We compared the marginal likelihoods of the full and reduced model (obtained through bridge sampling). This yielded a Bayes factor of 1.93 × 10^7^, indicating “decisive support” for the full model on Jeffreys’ scale.

## Discussion

We measured the impact of RIF resistance mutations on growth traits (maximum growth rate and maximum optical density in monoculture) in three species (*E. coli*, *A. baylyi* and *L. plantarum*) and two temperatures (30 *^◦^*C and 37 *^◦^*C). Our results indicate strong GxG effects between mutation and genetic background (including sign epistasis) as well as strong GxE and GxGxE effects that characterise this ‘fitness seascape’.

RIF resistance mutations in *rpoB* have been identified and characterised phenotypically in many species, including *Mycobacterium* spp., *E. coli*, *Bacillus* spp. and many others (e.g. Jin and Gross 1988; Vogler et al. 2002; Gagneux et al. 2006; Trindade et al. 2009; Goldstein 2014; Leehan and Nicholson 2021; Yang et al. 2023; Barilar et al. 2024; Bolourchi et al. 2025). It is clear from this large body of work that RIF resistance mutations themselves are generally highly conserved. However, to our knowledge only one previous study has directly compared fitness effects of RIF resistance mutations in the absence of RIF across species (Vogwill et al. 2016). Here, the authors measured the competitive fitness effects of homologous RIF resistance mutations in seven *Pseudomonas* species, reporting that those effects varied considerably across species: almost 40% of the variance in fitness was explained by interactions between mutations and species, and very little by mutation and species as individual effects. Even within a single species, the fitness effects of *rpoB* mutations can be highly variable, as shown by Hinz et al. (2024) for identical resistance mutations (conferring resistance to rifampicin, ciprofloxacin and streptomycin) across different clinical isolates of *E. coli*. Similarly, Trauner et al. (2021) showed that fitness effects of the S531L mutation vary markedly in different strains of *M. tuberculosis*. Knopp and Andersson (2018) appear to contradict these results, reporting predictable growth phenotypes for 13 chromosomal resistance mutations to different antibiotics across ten strains of *Salmonella enterica* and *E. coli* MG1655. However, the single RIF resistance mutation in their panel (S531L) did in fact exhibit considerable variation in its relative fitness effects in this study, even between strains that were almost identical. Overall, our results of strong epistatic effects across species are therefore not surprising, given that the three species we studied are only very distantly related (*E. coli* and *A. baylyi* are both gammaproteobacteria but belong to different orders, whereas *L. plantarum* belongs to a different phylum).

One mutation that has been phenotyped for growth in our study (in all three species) and in several previous studies is H526Y. This mutation consistently reduced competitive fitness in *Pseudomonas* but to widely varying degrees, ranging from almost no reduction to almost 27%. Moreover, most of the variation in competitive growth was explained by neither mutation nor species but by epistatic interactions between mutation and species, which in turn were not well explained by phylogenetic distance either (Vogwill et al. 2016). In clinical isolates of *E. coli*, H526Y had a wide range of effects on competitive fitness, with mostly negative effects but in some cases also increasing fitness (Hinz et al. 2024). In *M. tuberculosis*, H526Y causes a fitness reduction of around 20% (Gagneux et al. 2006). Broadly consistent with these results, H526Y reduced maximum growth rate in all three species that we investigated, but only weakly in *E. coli* and *L. plantarum* and without any epistatic interactions between those two species. By contrast, in *A. baylyi*, H526Y induced a reduction of around 16%, with consequently strong epistatic interactions with both other species.

Prior work has also established strong genotype-by-temperature interactions of RIF resistance mutations. Early experiments by Jin and Gross (1989) showed that some mutations in *rpoB* that confer RIF resistance also increased sensitivity to high and/or low temperatures in *E. coli*. Assuming that 37 *^◦^*C constitutes a mild heat stress in *A. baylyi*, this is in accord with our results that three mutants in this species had very low growth rates and reached very low maximum OD. However, at the level of individual mutation this parallelism does not hold: *E. coli* carrying mutation H526Y were unable to grow at 44 *^◦^*C in Jin and Gross (1989), but *A. baylyi* carrying the homologous mutation did not exhibit any particular heat sensitivity. This is also in line with experiments in *Pseudomonas aeruginosa* in which competitive growth of a H526Y mutant was not affected by high temperature (Gifford et al. 2016). Interestingly though, one of the three mutations that did induce heat sensitivity in *A. baylyi* in our experiments, H526D, is at the same site as H526Y; unfortunately H526D was not phenotyped in either of these two studies. Another mutation that has previously been reported to induce heat sensitivity, in *Listeria monocytogenes*, was D516G (Morse et al. 1999). Again, the corresponding *A. baylyi* mutant grew very well at 37 *^◦^*C in our experiments.

In addition to inducing heat-sensitivity, RIF resistance mutations in *rpoB* can also have the opposite effect, i.e. confer resistance to thermal stress. In *E. coli* populations that had been experimentally evolved for 2000 generations at high temperature (42.2 *^◦^*C) but in the absence of RIF, Rodŕıguez-Verdugo et al. (2013) observed the the repeated spread of four mutations that substantially increased competitive fitness at 42.2 *^◦^*C relative to the ancestor. Surprisingly though, these effects were very sensitive to genetic background: in a different *E. coli* strain (MG1655, the strain which we also studied), the same mutations reduced fitness at 42.2 *^◦^*C. One of these mutations (I572L) overlapped with our panel of mutants that we phenotyped in *A. baylyi* at 37 *^◦^*C, but this mutation affected maximum growth rate and maximum OD very little relative to the wildtype.

Similar, mutation-specific effects of temperature on fitness also seem to be involved in resistance to other antibiotics (Mira et al. 2022). Moreover, similar to temperature, strong GxE interac-tions have also be reported between RIF resistance mutations in *E. coli* and different nutrient conditions, including different growth media (Clarke et al. 2020; Soley et al. 2023; Hinz et al. 2024), different carbon sources (Gifford et al. 2016), and elemental nutrient limitation (Maharjan and Ferenci 2017). In many instances in these studies, not only the magnitude but also the direction of mutational fitness effects depended on the available nutrients, mirroring our and previous results on genotype by temperature interactions. Together, these results indicate that GxE interactions are pervasive in antibiotic resistance mutations.

How can we explain the wide range of GxG, GxE and GxGxE effects caused by single point mutations in the *rpoB* gene? Clearly, the RNAP and its beta subunit that *rpoB* codes for are central to the physiology of all bacteria, and as such pleiotropic effects of mutations in this gene that are unrelated to the binding affinity of RIF are to be expected. Mutations in *rpoB* have been shown to both increase or decrease the speed and efficiency of transcription (Qi et al. 2014; Yang et al. 2023). Mutations that increase the transcription speed (fast elongation rate and low pausing frequency) were reported to also rapidly deplete nucleotides within the cell, which in turn made them more sensitive to compounds such as 5FU and the antibiotic trimethoprim. Those mutants also had a growth advantage at low temperature (25 *^◦^*C) but grew slowly than the wildtype under heat stress (42 *^◦^*C) (Yang et al. 2023). In turn, differences in elongation rate and other properties of the RNAP appear to have global effects on gene expression within the cell (Qi et al. 2014; Rodŕıguez-Verdugo et al. 2016; Trauner et al. 2021; Soley et al. 2023). This transcriptional dysregulation has been shown to be highly idiosyncratic, with individual RIF resistance mutations in *rpoB* up- or down-regulating many genes in different ways (Qi et al. 2014; Trauner et al. 2021; Soley et al. 2023). Given that temperature also has large effects on gene expression, mutation by temperature interactions are expected to ensue. Interestingly, some RIF resistance mutations have been reported to largely reverse transcriptional changes brought about by heat stress (Rodŕıguez-Verdugo et al. 2016). Overall, although we are only beginning to understand the mechanistic basis of fitness effects of RIF resistance mutations, the findings of mutation-specific transcriptional dysregulation that is impacted by temperature provides at least some clues.

In conclusion, our study adds to the growing evidence indicating that cross-species predictions of fitness effects of antibiotic resistance mutations is extremely challenging, due to strong epistatic interactions between individual resistance mutations and genomic background. These problems are further compounded by strong environmental effects, in our case temperature, that give rise to both GxE and GxGxE interactions. As a result, accurate predictions of fitness effects of RIF resistance mutations, based on growth data in other species or other environments alone, appear impossible. It remains to be seen whether mechanistic models that integrate more data and processes—e.g. the effects of *rpoB* mutations on RNAP structure and enzymatic efficiency—can pave a way forward.

## Code availability

Code and data for this project will be made available on Github following publication.

## Acknowledgements

This research was supported by grants from the Australian Research Council (grants DP190102485 to JE and DP220103350 to ADL).

## Supplementary Material

**Figure S1:**
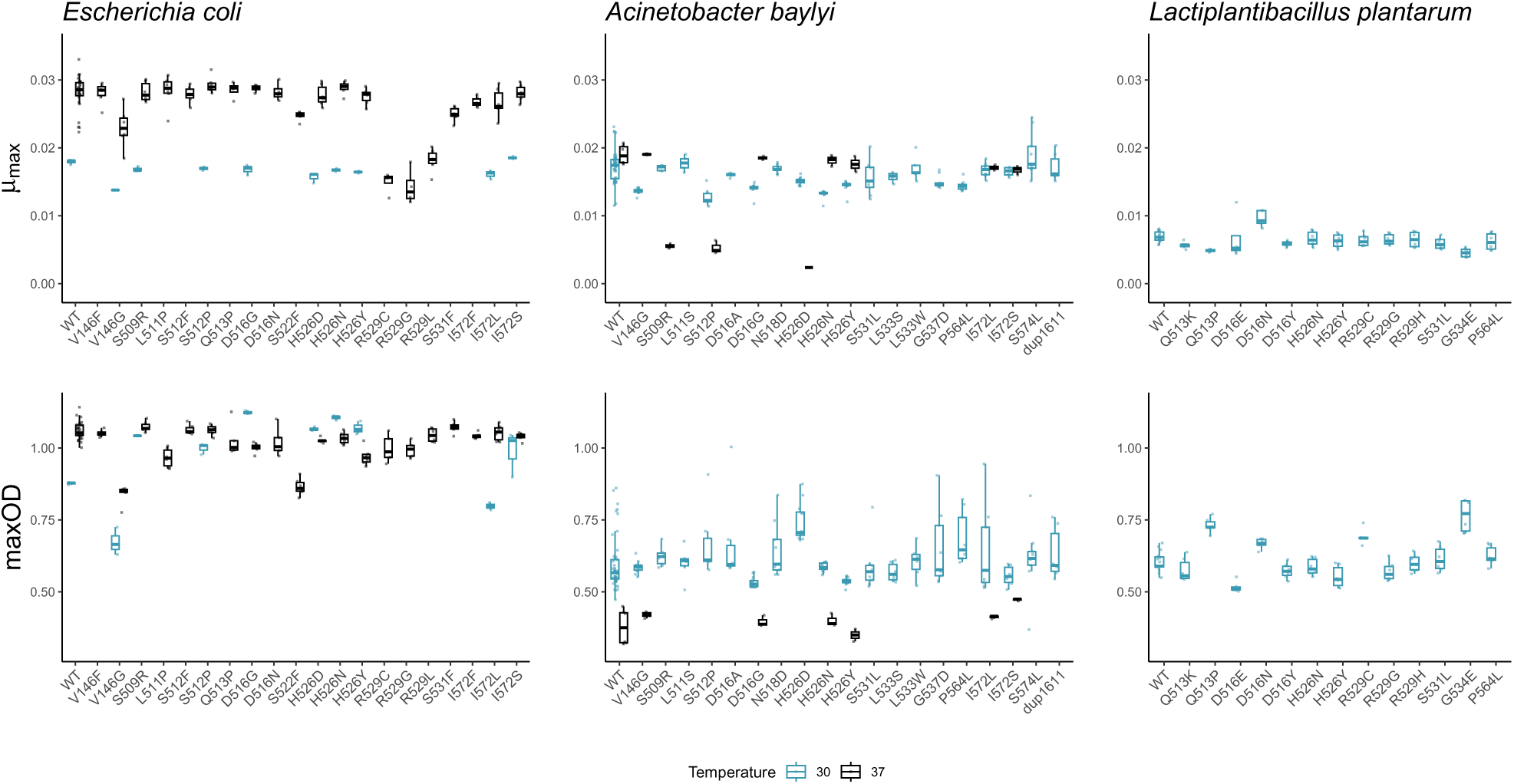
Measured growth traits for all three species, the two temperature conditions and all rifampicin resistant mutants. Small dots indicate estimates from individual growth assays and boxes show the median and interquartile range, with whiskers indicating the highest and lowest value within 1.5 × the interquartile range.

**Table S1:** Summary of the posterior distribution for the GxE model for mumax as the response variable, and mutation and temperature as independent variables. The columns show the mean, standard deviation, and credible interval of the posterior distribution for each estimated parameter. This model fit is for species *E. coli*. Formula used for fit: *µ*_max_ ∼ Mutation ∗ Temperature.

|  | Estimate | Est.Error | Q2.5 | Q97.5 |
| --- | --- | --- | --- | --- |
| b_Intercept | 0.0281 | 0.0003 | 0.0275 | 0.0288 |
| b_MutationD516G | 0.0006 | 0.0008 | -0.0009 | 0.0022 |
| b_MutationH526D | -0.0004 | 0.0008 | -0.0020 | 0.0013 |
| b_MutationH526N | 0.0008 | 0.0008 | -0.0008 | 0.0024 |
| b_MutationH526Y | -0.0006 | 0.0008 | -0.0021 | 0.0010 |
| b_MutationI572L | -0.0015 | 0.0008 | -0.0031 | 0.0001 |
| b_MutationI572S | -0.0000 | 0.0008 | -0.0016 | 0.0016 |
| b_MutationS509R | 0.0001 | 0.0008 | -0.0015 | 0.0017 |
| b_MutationS512P | 0.0011 | 0.0008 | -0.0005 | 0.0027 |
| b_MutationV146G | -0.0052 | 0.0008 | -0.0067 | -0.0035 |
| b_Temperature30 | -0.0102 | 0.0011 | -0.0123 | -0.0080 |
| b_MutationD516G:Temperature30 | -0.0018 | 0.0017 | -0.0050 | 0.0015 |
| b_MutationH526D:Temperature30 | -0.0019 | 0.0017 | -0.0054 | 0.0014 |
| b_MutationH526N:Temperature30 | -0.0020 | 0.0017 | -0.0052 | 0.0013 |
| b_MutationH526Y:Temperature30 | -0.0010 | 0.0017 | -0.0042 | 0.0024 |
| b_MutationI572L:Temperature30 | -0.0004 | 0.0017 | -0.0037 | 0.0028 |
| b_MutationI572S:Temperature30 | 0.0006 | 0.0017 | -0.0028 | 0.0039 |
| b_MutationS509R:Temperature30 | -0.0012 | 0.0017 | -0.0047 | 0.0021 |
| b_MutationS512P:Temperature30 | -0.0021 | 0.0017 | -0.0054 | 0.0011 |
| b_MutationV146G:Temperature30 | 0.0010 | 0.0017 | -0.0024 | 0.0044 |
| sigma | 0.0018 | 0.0001 | 0.0016 | 0.0021 |
| Intercept | 0.0248 | 0.0002 | 0.0244 | 0.0251 |
| lprior | -3.1413 | 0.0000 | -3.1413 | -3.1413 |
| lp_ | 540.1453 | 3.5393 | 532.2623 | 546.1706 |

**Table S2:**
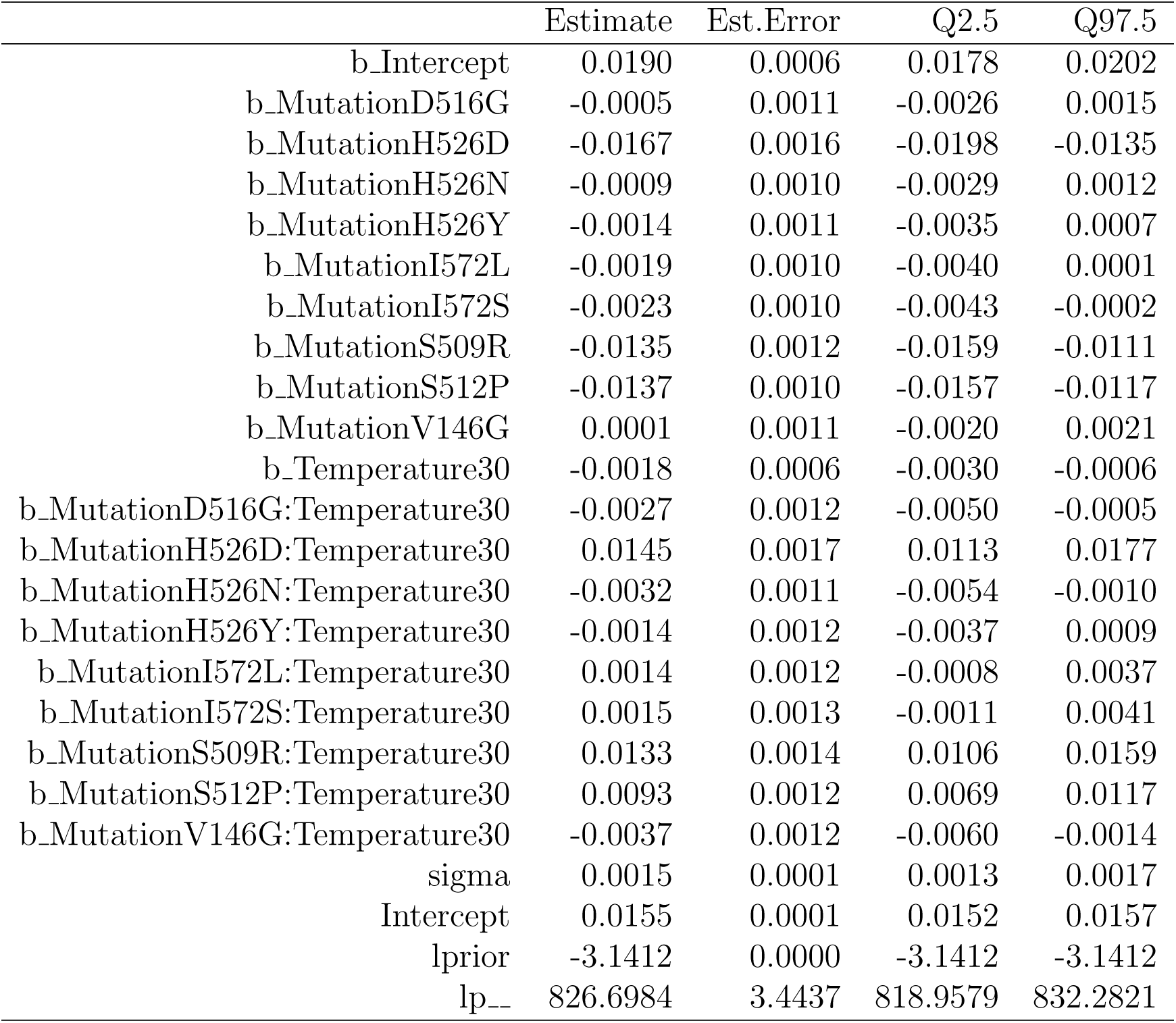
Summary of the posterior distribution for the GxE model for mumax as the response variable, and mutation and temperature as independent variables. The columns show the mean, standard deviation, and credible interval of the posterior distribution for each estimated parameter. This model fit is for species *A. baylyi*. Formula used for fit: *µ*_max_ ∼ Mutation ∗ Temperature.

**Table S3:** Summary of the posterior distribution for the GxE model for maxOD as the response variable, and mutation and temperature as independent variables. The columns show the mean, standard deviation, and credible interval of the posterior distribution for each estimated parameter. This model fit is for species *E. coli*. Formula used for fit: maxOD ∼ Mutation ∗ Temperature.

|  | Estimate | Est.Error | Q2.5 | Q97.5 |
| --- | --- | --- | --- | --- |
| b_Intercept | 1.0599 | 0.0052 | 1.0498 | 1.0704 |
| b_MutationD516G | -0.0587 | 0.0123 | -0.0821 | -0.0345 |
| b_MutationH526D | -0.0333 | 0.0125 | -0.0573 | -0.0091 |
| b_MutationH526N | -0.0257 | 0.0125 | -0.0497 | -0.0009 |
| b_MutationH526Y | -0.0893 | 0.0123 | -0.1132 | -0.0648 |
| b_MutationI572L | -0.0071 | 0.0123 | -0.0320 | 0.0176 |
| b_MutationI572S | -0.0207 | 0.0122 | -0.0451 | 0.0033 |
| b_MutationS509R | 0.0141 | 0.0123 | -0.0105 | 0.0379 |
| b_MutationS512P | 0.0027 | 0.0125 | -0.0214 | 0.0274 |
| b_MutationV146G | -0.2204 | 0.0127 | -0.2467 | -0.1966 |
| b_Temperature30 | -0.1832 | 0.0164 | -0.2147 | -0.1515 |
| b_MutationD516G:Temperature30 | 0.3061 | 0.0248 | 0.2573 | 0.3536 |
| b_MutationH526D:Temperature30 | 0.2226 | 0.0252 | 0.1709 | 0.2717 |
| b_MutationH526N:Temperature30 | 0.2549 | 0.0252 | 0.2051 | 0.3031 |
| b_MutationH526Y:Temperature30 | 0.2820 | 0.0256 | 0.2310 | 0.3314 |
| b_MutationI572L:Temperature30 | -0.0713 | 0.0250 | -0.1207 | -0.0230 |
| b_MutationI572S:Temperature30 | 0.1340 | 0.0252 | 0.0841 | 0.1849 |
| b_MutationS509R:Temperature30 | 0.1515 | 0.0248 | 0.1029 | 0.1998 |
| b_MutationS512P:Temperature30 | 0.1198 | 0.0256 | 0.0701 | 0.1704 |
| b_MutationV146G:Temperature30 | 0.0171 | 0.0255 | -0.0327 | 0.0678 |
| sigma | 0.0274 | 0.0021 | 0.0236 | 0.0318 |
| Intercept | 1.0140 | 0.0027 | 1.0087 | 1.0191 |
| lprior | -3.1413 | 0.0000 | -3.1413 | -3.1413 |
| lp__ | 240.3367 | 3.4888 | 232.5543 | 246.1141 |

**Table S4:** Summary of the posterior distribution for the GxE model for maxOD as the response variable, and mutation and temperature as independent variables. The columns show the mean, standard deviation, and credible interval of the posterior distribution for each estimated parameter. This model fit is for species *A. baylyi*. Formula used for fit: maxOD ∼ Mutation ∗ Temperature.

|  | Estimate | Est.Error | Q2.5 | Q97.5 |
| --- | --- | --- | --- | --- |
| b_Intercept | 0.3784 | 0.0295 | 0.3222 | 0.4355 |
| b_MutationD516G | 0.0181 | 0.0502 | -0.0835 | 0.1169 |
| b_MutationH526D | -0.3734 | 0.0462 | -0.4654 | -0.2849 |
| b_MutationH526N | 0.0222 | 0.0501 | -0.0770 | 0.1199 |
| b_MutationH526Y | -0.0294 | 0.0498 | -0.1252 | 0.0678 |
| b_MutationI572L | 0.0351 | 0.0506 | -0.0621 | 0.1338 |
| b_MutationI572S | 0.0938 | 0.0485 | -0.0045 | 0.1876 |
| b_MutationS509R | -0.3538 | 0.0574 | -0.4671 | -0.2400 |
| b_MutationS512P | -0.3044 | 0.0503 | -0.4044 | -0.2070 |
| b_MutationV146G | 0.0428 | 0.0503 | -0.0553 | 0.1418 |
| b_Temperature30 | 0.2175 | 0.0312 | 0.1567 | 0.2779 |
| b_MutationD516G:Temperature30 | -0.0815 | 0.0543 | -0.1884 | 0.0294 |
| b_MutationH526D:Temperature30 | 0.5200 | 0.0510 | 0.4223 | 0.6193 |
| b_MutationH526N:Temperature30 | -0.0311 | 0.0550 | -0.1366 | 0.0765 |
| b_MutationH526Y:Temperature30 | -0.0300 | 0.0540 | -0.1353 | 0.0749 |
| b_MutationI572L:Temperature30 | 0.0016 | 0.0564 | -0.1075 | 0.1078 |
| b_MutationI572S:Temperature30 | -0.1340 | 0.0570 | -0.2474 | -0.0210 |
| b_MutationS509R:Temperature30 | 0.3817 | 0.0656 | 0.2562 | 0.5084 |
| b_MutationS512P:Temperature30 | 0.3797 | 0.0584 | 0.2664 | 0.4932 |
| b_MutationV146G:Temperature30 | -0.0527 | 0.0548 | -0.1594 | 0.0544 |
| sigma | 0.0700 | 0.0042 | 0.0625 | 0.0789 |
| Intercept | 0.5436 | 0.0053 | 0.5333 | 0.5540 |
| lprior | -3.1421 | 0.0001 | -3.1423 | -3.1419 |
| lp__ | 214.2521 | 3.5621 | 206.2013 | 220.0754 |

**Table S5:** Summary of the posterior distribution for the GxG model for mumax as the response variable, and mutation and species as independent variables. The columns show the mean, standard deviation, and credible interval of the posterior distribution for each estimated parameter. This model fit is for species pair *E. coli* / *A. baylyi*. Formula used for fit: *µ*_max_ ∼ Mutation ∗ Species.

|  | Estimate | Est.Error | Q2.5 | Q97.5 |
| --- | --- | --- | --- | --- |
| b_Intercept | 0.0281 | 0.0003 | 0.0274 | 0.0288 |
| b_MutationD516G | 0.0006 | 0.0008 | -0.0009 | 0.0022 |
| b_MutationH526D | -0.0004 | 0.0008 | -0.0019 | 0.0012 |
| b_MutationH526N | 0.0008 | 0.0008 | -0.0007 | 0.0023 |
| b_MutationH526Y | -0.0005 | 0.0008 | -0.0020 | 0.0010 |
| b_MutationI572L | -0.0015 | 0.0008 | -0.0031 | 0.0002 |
| b_MutationI572S | -0.0000 | 0.0008 | -0.0015 | 0.0016 |
| b_MutationS509R | 0.0001 | 0.0008 | -0.0015 | 0.0016 |
| b_MutationS512P | 0.0012 | 0.0008 | -0.0004 | 0.0027 |
| b_MutationV146G | -0.0051 | 0.0008 | -0.0067 | -0.0036 |
| b_SpeciesAbaylyi | -0.0109 | 0.0004 | -0.0117 | -0.0100 |
| b_MutationD516G:SpeciesAbaylyi | -0.0039 | 0.0009 | -0.0057 | -0.0020 |
| b_MutationH526D:SpeciesAbaylyi | -0.0018 | 0.0010 | -0.0037 | 0.0002 |
| b_MutationH526N:SpeciesAbaylyi | -0.0048 | 0.0010 | -0.0067 | -0.0029 |
| b_MutationH526Y:SpeciesAbaylyi | -0.0023 | 0.0009 | -0.0041 | -0.0005 |
| b_MutationI572L:SpeciesAbaylyi | 0.0010 | 0.0010 | -0.0011 | 0.0030 |
| b_MutationI572S:SpeciesAbaylyi | -0.0008 | 0.0012 | -0.0032 | 0.0016 |
| b_MutationS509R:SpeciesAbaylyi | -0.0003 | 0.0011 | -0.0025 | 0.0020 |
| b_MutationS512P:SpeciesAbaylyi | -0.0056 | 0.0011 | -0.0078 | -0.0034 |
| b_MutationV146G:SpeciesAbaylyi | 0.0015 | 0.0010 | -0.0003 | 0.0034 |
| sigma | 0.0017 | 0.0001 | 0.0016 | 0.0019 |
| Intercept | 0.0201 | 0.0001 | 0.0199 | 0.0204 |
| lprior | -3.1413 | 0.0000 | -3.1413 | -3.1413 |
| lp_ | 1058.2413 | 3.4551 | 1050.6565 | 1063.8246 |

**Table S6:** Summary of the posterior distribution for the GxG model for mumax as the response variable, and mutation and species as independent variables. The columns show the mean, standard deviation, and credible interval of the posterior distribution for each estimated parameter. This model fit is for species pair *E. coli* / *L. plantarum*. Formula used for fit: *µ*_max_ ∼ Mutation∗Species.

|  | Estimate | Est.Error | Q2.5 | Q97.5 |
| --- | --- | --- | --- | --- |
| b_Intercept | 0.0281 | 0.0003 | 0.0275 | 0.0287 |
| b_MutationD516N | 0.0001 | 0.0009 | -0.0017 | 0.0018 |
| b_MutationH526N | 0.0007 | 0.0007 | -0.0008 | 0.0022 |
| b_MutationH526Y | -0.0005 | 0.0008 | -0.0020 | 0.0009 |
| b_MutationQ513P | 0.0004 | 0.0009 | -0.0013 | 0.0022 |
| b_MutationR529C | -0.0132 | 0.0009 | -0.0149 | -0.0115 |
| b_MutationR529G | -0.0139 | 0.0009 | -0.0156 | -0.0121 |
| b_SpeciesLplantarum | -0.0212 | 0.0005 | -0.0222 | -0.0202 |
| b_MutationD516N:SpeciesLplantarum | 0.0025 | 0.0013 | 0.0001 | 0.0051 |
| b_MutationH526N:SpeciesLplantarum | -0.0010 | 0.0011 | -0.0031 | 0.0011 |
| b_MutationH526Y:SpeciesLplantarum | -0.0001 | 0.0011 | -0.0023 | 0.0021 |
| b_MutationQ513P:SpeciesLplantarum | -0.0025 | 0.0012 | -0.0048 | -0.0002 |
| b_MutationR529C:SpeciesLplantarum | 0.0126 | 0.0012 | 0.0103 | 0.0150 |
| b_MutationR529G:SpeciesLplantarum | 0.0135 | 0.0012 | 0.0111 | 0.0157 |
| sigma | 0.0017 | 0.0001 | 0.0014 | 0.0019 |
| Intercept | 0.0170 | 0.0002 | 0.0167 | 0.0173 |
| lprior | -3.1412 | 0.0000 | -3.1412 | -3.1412 |
| lp_ | 518.3420 | 2.8872 | 511.7350 | 522.9177 |

**Table S7:** Summary of the posterior distribution for the GxG model for mumax as the response variable, and mutation and species as independent variables. The columns show the mean, standard deviation, and credible interval of the posterior distribution for each estimated parameter. This model fit is for species pair *A. baylyi* / *L. plantarum*. Formula used for fit: *µ*_max_ ∼ Mutation ∗ Species.

|  | Estimate | Est.Error | Q2.5 | Q97.5 |
| --- | --- | --- | --- | --- |
| b_Intercept | 0.0172 | 0.0003 | 0.0167 | 0.0177 |
| b_MutationH526N | -0.0040 | 0.0005 | -0.0051 | -0.0030 |
| b_MutationH526Y | -0.0028 | 0.0005 | -0.0039 | -0.0018 |
| b_MutationP564L | -0.0028 | 0.0006 | -0.0040 | -0.0015 |
| b_MutationS531L | -0.0016 | 0.0007 | -0.0029 | -0.0003 |
| b_SpeciesLplantarum | -0.0103 | 0.0005 | -0.0113 | -0.0093 |
| b_MutationH526N:SpeciesLplantarum | 0.0037 | 0.0010 | 0.0018 | 0.0055 |
| b_MutationH526Y:SpeciesLplantarum | 0.0022 | 0.0010 | 0.0002 | 0.0042 |
| b_MutationP564L:SpeciesLplantarum | 0.0021 | 0.0010 | 0.0000 | 0.0041 |
| b_MutationS531L:SpeciesLplantarum | 0.0006 | 0.0011 | -0.0015 | 0.0027 |
| sigma | 0.0017 | 0.0001 | 0.0015 | 0.0020 |
| Intercept | 0.0130 | 0.0002 | 0.0127 | 0.0133 |
| lprior | -3.1412 | 0.0000 | -3.1412 | -3.1412 |
| lp_ | 624.0497 | 2.5026 | 617.9604 | 627.8308 |

**Table S8:**
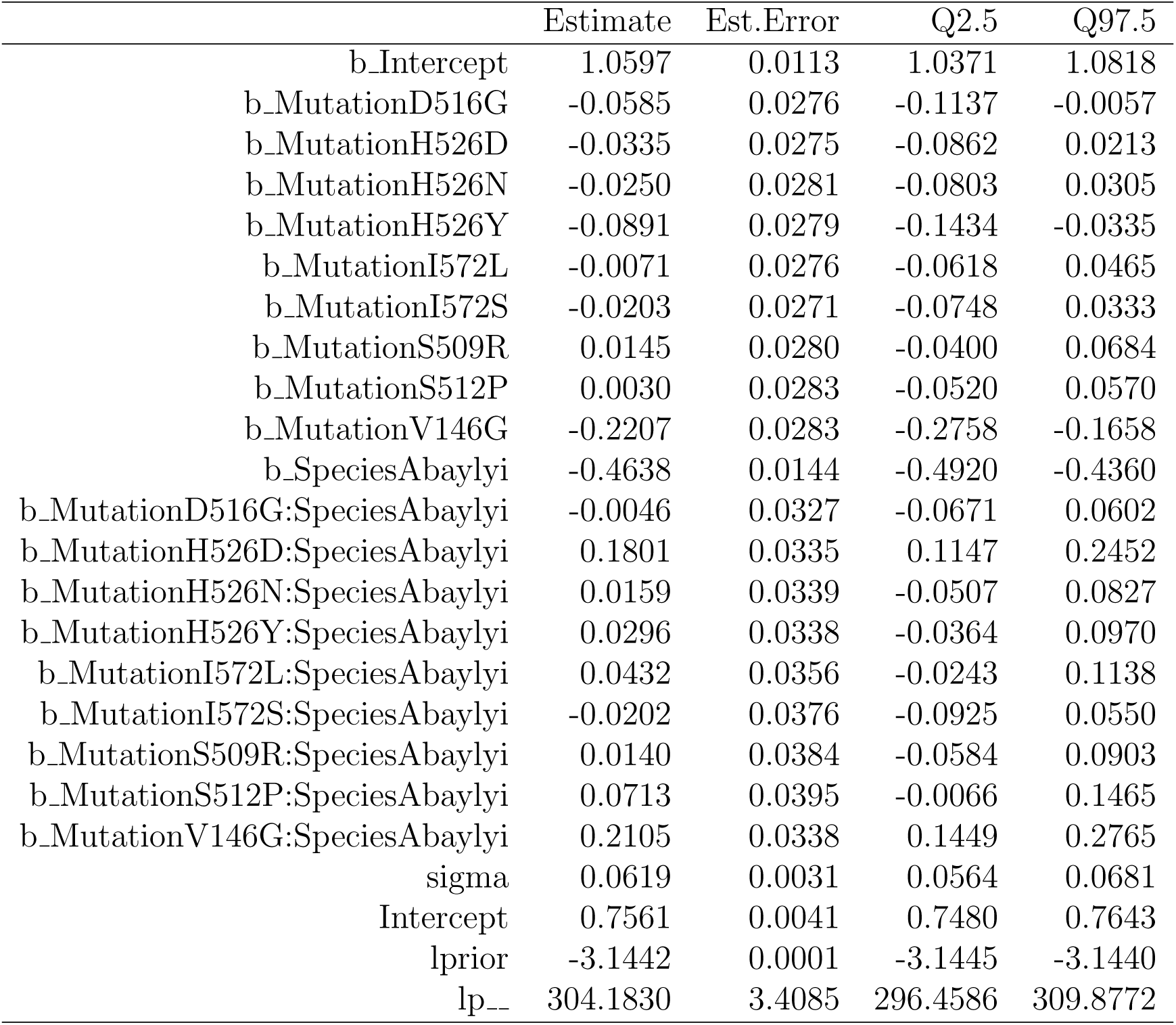
Summary of the posterior distribution for the GxG model for maxOD as the response variable, and mutation and species as independent variables. The columns show the mean, standard deviation, and credible interval of the posterior distribution for each estimated parameter. This model fit is for species pair *E. coli* / *A. baylyi*. Formula used for fit: maxOD ∼ Mutation ∗ Species.

|  | Estimate | Est.Error | Q2.5 | Q97.5 |
| --- | --- | --- | --- | --- |
| b_Intercept | 1.0597 | 0.0113 | 1.0371 | 1.0818 |
| b_MutationD516G | -0.0585 | 0.0276 | -0.1137 | -0.0057 |
| b_MutationH526D | -0.0335 | 0.0275 | -0.0862 | 0.0213 |
| b_MutationH526N | -0.0250 | 0.0281 | -0.0803 | 0.0305 |
| b_MutationH526Y | -0.0891 | 0.0279 | -0.1434 | -0.0335 |
| b_MutationI572L | -0.0071 | 0.0276 | -0.0618 | 0.0465 |
| b_MutationI572S | -0.0203 | 0.0271 | -0.0748 | 0.0333 |
| b_MutationS509R | 0.0145 | 0.0280 | -0.0400 | 0.0684 |
| b_MutationS512P | 0.0030 | 0.0283 | -0.0520 | 0.0570 |
| b_MutationV146G | -0.2207 | 0.0283 | -0.2758 | -0.1658 |
| b_SpeciesAbaylyi | -0.4638 | 0.0144 | -0.4920 | -0.4360 |
| b_MutationD516G:SpeciesAbaylyi | -0.0046 | 0.0327 | -0.0671 | 0.0602 |
| b_MutationH526D:SpeciesAbaylyi | 0.1801 | 0.0335 | 0.1147 | 0.2452 |
| b_MutationH526N:SpeciesAbaylyi | 0.0159 | 0.0339 | -0.0507 | 0.0827 |
| b_MutationH526Y:SpeciesAbaylyi | 0.0296 | 0.0338 | -0.0364 | 0.0970 |
| b_MutationI572L:SpeciesAbaylyi | 0.0432 | 0.0356 | -0.0243 | 0.1138 |
| b_MutationI572S:SpeciesAbaylyi | -0.0202 | 0.0376 | -0.0925 | 0.0550 |
| b_MutationS509R:SpeciesAbaylyi | 0.0140 | 0.0384 | -0.0584 | 0.0903 |
| b_MutationS512P:SpeciesAbaylyi | 0.0713 | 0.0395 | -0.0066 | 0.1465 |
| b_MutationV146G:SpeciesAbaylyi | 0.2105 | 0.0338 | 0.1449 | 0.2765 |
| sigma | 0.0619 | 0.0031 | 0.0564 | 0.0681 |
| Intercept | 0.7561 | 0.0041 | 0.7480 | 0.7643 |
| lprior | -3.1442 | 0.0001 | -3.1445 | -3.1440 |
| lp__ | 304.1830 | 3.4085 | 296.4586 | 309.8772 |

**Table S9:** Summary of the posterior distribution for the GxG model for maxOD as the response variable, and mutation and species as independent variables. The columns show the mean, standard deviation, and credible interval of the posterior distribution for each estimated parameter. This model fit is for species pair *E. coli* / *L. plantarum*. Formula used for fit: maxOD ∼ Mutation ∗ Species.

|  | Estimate | Est.Error | Q2.5 | Q97.5 |
| --- | --- | --- | --- | --- |
| b_Intercept | 1.0598 | 0.0064 | 1.0474 | 1.0724 |
| b_MutationD516N | -0.0383 | 0.0166 | -0.0702 | -0.0058 |
| b_MutationH526N | -0.0255 | 0.0154 | -0.0554 | 0.0044 |
| b_MutationH526Y | -0.0892 | 0.0155 | -0.1188 | -0.0570 |
| b_MutationQ513P | -0.0325 | 0.0165 | -0.0650 | 0.0001 |
| b_MutationR529C | -0.0607 | 0.0166 | -0.0925 | -0.0277 |
| b_MutationR529G | -0.0651 | 0.0169 | -0.0977 | -0.0318 |
| b_SpeciesLplantarum | -0.4554 | 0.0109 | -0.4768 | -0.4340 |
| b_MutationD516N:SpeciesLplantarum | 0.1025 | 0.0234 | 0.0559 | 0.1490 |
| b_MutationH526N:SpeciesLplantarum | 0.0078 | 0.0225 | -0.0366 | 0.0512 |
| b_MutationH526Y:SpeciesLplantarum | 0.0368 | 0.0226 | -0.0093 | 0.0800 |
| b_MutationQ513P:SpeciesLplantarum | 0.1590 | 0.0230 | 0.1142 | 0.2049 |
| b_MutationR529C:SpeciesLplantarum | 0.1478 | 0.0237 | 0.1008 | 0.1937 |
| b_MutationR529G:SpeciesLplantarum | 0.0317 | 0.0239 | -0.0150 | 0.0783 |
| sigma | 0.0340 | 0.0025 | 0.0296 | 0.0392 |
| Intercept | 0.8469 | 0.0031 | 0.8406 | 0.8530 |
| lprior | -3.1438 | 0.0001 | -3.1440 | -3.1436 |
| lp_-- | 213.9886 | 2.8610 | 207.7304 | 218.7550 |

**Table S10:** Summary of the posterior distribution for the GxG model for maxOD as the response variable, and mutation and species as independent variables. The columns show the mean, standard deviation, and credible interval of the posterior distribution for each estimated parameter. This model fit is for species pair *A. baylyi* / *L. plantarum*. Formula used for fit: maxOD ∼ Mutation ∗ Species.

|  | Estimate | Est.Error | Q2.5 | Q97.5 |
| --- | --- | --- | --- | --- |
| b_Intercept | 0.5958 | 0.0096 | 0.5770 | 0.6144 |
| b_MutationH526N | -0.0087 | 0.0206 | -0.0488 | 0.0327 |
| b_MutationH526Y | -0.0596 | 0.0203 | -0.0990 | -0.0205 |
| b_MutationP564L | 0.0886 | 0.0245 | 0.0419 | 0.1371 |
| b_MutationS531L | -0.0104 | 0.0248 | -0.0595 | 0.0379 |
| b_SpeciesLplantarum | 0.0087 | 0.0200 | -0.0317 | 0.0474 |
| b_MutationH526N:SpeciesLplantarum | -0.0094 | 0.0384 | -0.0870 | 0.0678 |
| b_MutationH526Y:SpeciesLplantarum | 0.0072 | 0.0394 | -0.0688 | 0.0852 |
| b_MutationP564L:SpeciesLplantarum | -0.0671 | 0.0405 | -0.1451 | 0.0123 |
| b_MutationS531L:SpeciesLplantarum | 0.0202 | 0.0422 | -0.0651 | 0.1004 |
| sigma | 0.0687 | 0.0044 | 0.0608 | 0.0782 |
| Intercept | 0.5947 | 0.0059 | 0.5831 | 0.6062 |
| lprior | -3.1417 | 0.0001 | -3.1419 | -3.1416 |
| lp_ | 163.4796 | 2.5057 | 157.4632 | 167.2444 |

**Table S11:** Summary of the posterior distribution for the GxGxE model for mumax as the response variable, and mutation, temperature and species as independent variables. The columns show the mean, standard deviation, and credible interval of the posterior distribution for each estimated parameter. This model fit is for species pair *E. coli* / *A. baylyi*. Formula used for fit: *µ*_max_ ∼ Mutation ∗ Species ∗ Temperature.

|  | Estimate | Est.Error | Q2.5 | Q97.5 |
| --- | --- | --- | --- | --- |
| b_Intercept | 0.0281 | 0.0003 | 0.0275 | 0.0287 |
| b_MutationD516G | 0.0007 | 0.0007 | -0.0007 | 0.0020 |
| b_MutationH526D | -0.0004 | 0.0007 | -0.0018 | 0.0010 |
| b_MutationH526N | 0.0008 | 0.0007 | -0.0006 | 0.0021 |
| b_MutationH526Y | -0.0006 | 0.0007 | -0.0020 | 0.0008 |
| b_MutationI572L | -0.0015 | 0.0007 | -0.0029 | -0.0001 |
| b_MutationI572S | -0.0000 | 0.0007 | -0.0014 | 0.0014 |
| b_MutationS509R | 0.0001 | 0.0007 | -0.0013 | 0.0015 |
| b_MutationS512P | 0.0011 | 0.0007 | -0.0003 | 0.0025 |
| b_MutationV146G | -0.0051 | 0.0007 | -0.0065 | -0.0037 |
| b_SpeciesAbaylyi | -0.0091 | 0.0007 | -0.0105 | -0.0077 |
| b_Temperature30 | -0.0102 | 0.0010 | -0.0121 | -0.0083 |
| b_MutationD516G:SpeciesAbaylyi | -0.0011 | 0.0013 | -0.0036 | 0.0015 |
| b_MutationH526D:SpeciesAbaylyi | -0.0162 | 0.0019 | -0.0198 | -0.0125 |
| b_MutationH526N:SpeciesAbaylyi | -0.0016 | 0.0013 | -0.0042 | 0.0011 |
| b_MutationH526Y:SpeciesAbaylyi | -0.0008 | 0.0014 | -0.0035 | 0.0018 |
| b_MutationI572L:SpeciesAbaylyi | -0.0004 | 0.0013 | -0.0031 | 0.0021 |
| b_MutationI572S:SpeciesAbaylyi | -0.0023 | 0.0013 | -0.0049 | 0.0003 |
| b_MutationS509R:SpeciesAbaylyi | -0.0135 | 0.0015 | -0.0163 | -0.0106 |
| b_MutationS512P:SpeciesAbaylyi | -0.0148 | 0.0013 | -0.0174 | -0.0122 |
| b_MutationV146G:SpeciesAbaylyi | 0.0052 | 0.0014 | 0.0026 | 0.0079 |
| b_MutationD516G:Temperature30 | -0.0018 | 0.0015 | -0.0047 | 0.0011 |
| b_MutationH526D:Temperature30 | -0.0019 | 0.0015 | -0.0048 | 0.0010 |
| b_MutationH526N:Temperature30 | -0.0020 | 0.0015 | -0.0050 | 0.0010 |
| b_MutationH526Y:Temperature30 | -0.0010 | 0.0015 | -0.0039 | 0.0020 |
| b_MutationI572L:Temperature30 | -0.0004 | 0.0014 | -0.0031 | 0.0025 |
| b_MutationI572S:Temperature30 | 0.0006 | 0.0015 | -0.0023 | 0.0035 |
| b_MutationS509R:Temperature30 | -0.0012 | 0.0015 | -0.0041 | 0.0016 |
| b_MutationS512P:Temperature30 | -0.0021 | 0.0015 | -0.0050 | 0.0008 |
| b_MutationV146G:Temperature30 | 0.0009 | 0.0015 | -0.0020 | 0.0039 |
| b_SpeciesAbaylyi:Temperature30 | 0.0084 | 0.0012 | 0.0061 | 0.0107 |
| b_MutationD516G:SpeciesAbaylyi:Temperature30 | -0.0009 | 0.0019 | -0.0047 | 0.0027 |
| b_MutationH526D:SpeciesAbaylyi:Temperature30 | 0.0164 | 0.0023 | 0.0117 | 0.0210 |
| b_MutationH526N:SpeciesAbaylyi:Temperature30 | -0.0012 | 0.0019 | -0.0050 | 0.0025 |
| b_MutationH526Y:SpeciesAbaylyi:Temperature30 | -0.0004 | 0.0019 | -0.0042 | 0.0032 |
| b_MutationI572L:SpeciesAbaylyi:Temperature30 | 0.0018 | 0.0019 | -0.0020 | 0.0055 |
| b_MutationI572S:SpeciesAbaylyi:Temperature30 | 0.0009 | 0.0020 | -0.0030 | 0.0048 |
| b_MutationS509R:SpeciesAbaylyi:Temperature30 | 0.0144 | 0.0021 | 0.0103 | 0.0185 |
| b_MutationS512P:SpeciesAbaylyi:Temperature30 | 0.0114 | 0.0020 | 0.0074 | 0.0152 |
| b_MutationV146G:SpeciesAbaylyi:Temperature30 | -0.0046 | 0.0019 | -0.0083 | -0.0009 |
| sigma | 0.0016 | 0.0001 | 0.0015 | 0.0018 |
| Intercept | 0.0192 | 0.0001 | 0.0190 | 0.0194 |
| lprior | -3.1413 | 0.0000 | -3.1413 | -3.1413 |
| lp_ | 1375.6707 | 4.7964 | 1365.0407 | 1383.9234 |

**Table S12:**
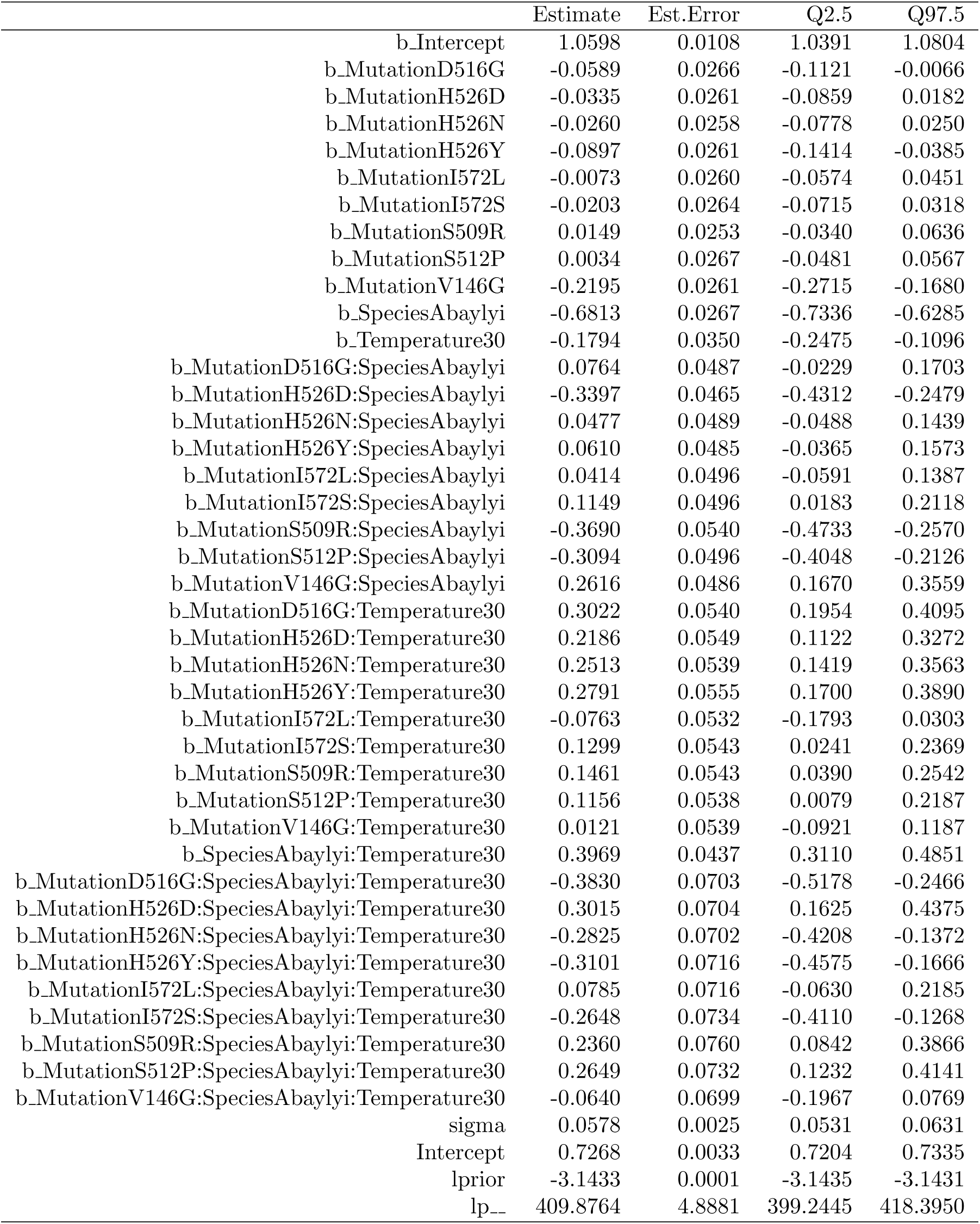
Summary of the posterior distribution for the GxGxE model for maxOD as the response variable, and mutation, temperature and species as independent variables. The columns show the mean, standard deviation, and credible interval of the posterior distribution for each estimated parameter. This model fit is for species pair *E. coli* / *A. baylyi*. Formula used for fit: maxOD ∼ Mutation ∗ Species ∗ Temperature.

**Table S13:**
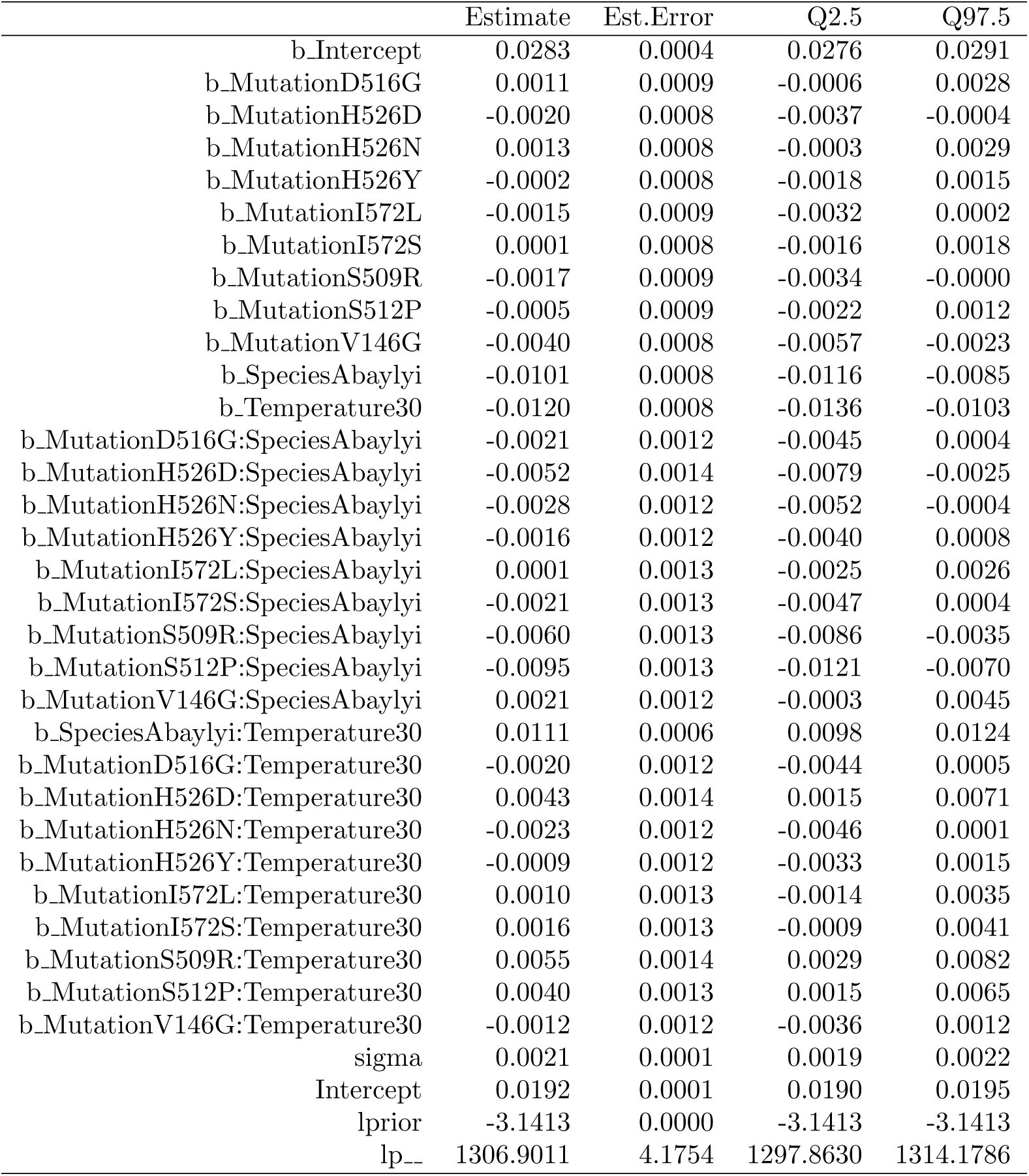
Summary of the posterior distribution for the GxGxE model for mumax as the response variable, and mutation, temperature and species as independent variables. The columns show the mean, standard deviation, and credible interval of the posterior distribution for each estimated parameter. This model fit is for species pair *E. coli* / *A. baylyi*. Formula used for fit: *µ*_max_ ∼ Mutation ∗ Species + Temperature ∗ Species + Mutation ∗ Temperature.

|  | Estimate | Est.Error | Q2.5 | Q97.5 |
| --- | --- | --- | --- | --- |
| b_Intercept | 0.0283 | 0.0004 | 0.0276 | 0.0291 |
| b_MutationD516G | 0.0011 | 0.0009 | -0.0006 | 0.0028 |
| b_MutationH526D | -0.0020 | 0.0008 | -0.0037 | -0.0004 |
| b_MutationH526N | 0.0013 | 0.0008 | -0.0003 | 0.0029 |
| b_MutationH526Y | -0.0002 | 0.0008 | -0.0018 | 0.0015 |
| b_MutationI572L | -0.0015 | 0.0009 | -0.0032 | 0.0002 |
| b_MutationI572S | 0.0001 | 0.0008 | -0.0016 | 0.0018 |
| b_MutationS509R | -0.0017 | 0.0009 | -0.0034 | -0.0000 |
| b_MutationS512P | -0.0005 | 0.0009 | -0.0022 | 0.0012 |
| b_MutationV146G | -0.0040 | 0.0008 | -0.0057 | -0.0023 |
| b_SpeciesAbaylyi | -0.0101 | 0.0008 | -0.0116 | -0.0085 |
| b_Temperature30 | -0.0120 | 0.0008 | -0.0136 | -0.0103 |
| b_MutationD516G:SpeciesAbaylyi | -0.0021 | 0.0012 | -0.0045 | 0.0004 |
| b_MutationH526D:SpeciesAbaylyi | -0.0052 | 0.0014 | -0.0079 | -0.0025 |
| b_MutationH526N:SpeciesAbaylyi | -0.0028 | 0.0012 | -0.0052 | -0.0004 |
| b_MutationH526Y:SpeciesAbaylyi | -0.0016 | 0.0012 | -0.0040 | 0.0008 |
| b_MutationI572L:SpeciesAbaylyi | 0.0001 | 0.0013 | -0.0025 | 0.0026 |
| b_MutationI572S:SpeciesAbaylyi | -0.0021 | 0.0013 | -0.0047 | 0.0004 |
| b_MutationS509R:SpeciesAbaylyi | -0.0060 | 0.0013 | -0.0086 | -0.0035 |
| b_MutationS512P:SpeciesAbaylyi | -0.0095 | 0.0013 | -0.0121 | -0.0070 |
| b_MutationV146G:SpeciesAbaylyi | 0.0021 | 0.0012 | -0.0003 | 0.0045 |
| b_SpeciesAbaylyi:Temperature30 | 0.0111 | 0.0006 | 0.0098 | 0.0124 |
| b_MutationD516G:Temperature30 | -0.0020 | 0.0012 | -0.0044 | 0.0005 |
| b_MutationH526D:Temperature30 | 0.0043 | 0.0014 | 0.0015 | 0.0071 |
| b_MutationH526N:Temperature30 | -0.0023 | 0.0012 | -0.0046 | 0.0001 |
| b_MutationH526Y:Temperature30 | -0.0009 | 0.0012 | -0.0033 | 0.0015 |
| b_MutationI572L:Temperature30 | 0.0010 | 0.0013 | -0.0014 | 0.0035 |
| b_MutationI572S:Temperature30 | 0.0016 | 0.0013 | -0.0009 | 0.0041 |
| b_MutationS509R:Temperature30 | 0.0055 | 0.0014 | 0.0029 | 0.0082 |
| b_MutationS512P:Temperature30 | 0.0040 | 0.0013 | 0.0015 | 0.0065 |
| b_MutationV146G:Temperature30 | -0.0012 | 0.0012 | -0.0036 | 0.0012 |
| sigma | 0.0021 | 0.0001 | 0.0019 | 0.0022 |
| Intercept | 0.0192 | 0.0001 | 0.0190 | 0.0195 |
| lprior | -3.1413 | 0.0000 | -3.1413 | -3.1413 |
| lp_ | 1306.9011 | 4.1754 | 1297.8630 | 1314.1786 |

**Table S14:** Summary of the posterior distribution for the GxGxE model for maxOD as the response variable, and mutation, temperature and species as independent variables. The columns show the mean, standard deviation, and credible interval of the posterior distribution for each estimated parameter. This model fit is for species pair *E. coli* / *A. baylyi*. Formula used for fit: maxOD ∼ Mutation ∗ Species + Temperature ∗ Species + Mutation ∗ Temperature.

|  | Estimate | Est.Error | Q2.5 | Q97.5 |
| --- | --- | --- | --- | --- |
| b_Intercept | 1.0569 | 0.0140 | 1.0290 | 1.0837 |
| b_MutationD516G | 0.0071 | 0.0313 | -0.0540 | 0.0681 |
| b_MutationH526D | -0.0991 | 0.0313 | -0.1624 | -0.0392 |
| b_MutationH526N | 0.0219 | 0.0310 | -0.0400 | 0.0829 |
| b_MutationH526Y | -0.0372 | 0.0317 | -0.0981 | 0.0245 |
| b_MutationI572L | -0.0251 | 0.0311 | -0.0857 | 0.0359 |
| b_MutationI572S | 0.0191 | 0.0321 | -0.0429 | 0.0834 |
| b_MutationS509R | -0.0224 | 0.0325 | -0.0858 | 0.0409 |
| b_MutationS512P | -0.0452 | 0.0318 | -0.1089 | 0.0174 |
| b_MutationV146G | -0.2124 | 0.0316 | -0.2736 | -0.1495 |
| b_SpeciesAbaylyi | -0.6644 | 0.0276 | -0.7183 | -0.6113 |
| b_Temperature30 | -0.1514 | 0.0306 | -0.2126 | -0.0914 |
| b_MutationD516G:SpeciesAbaylyi | -0.1283 | 0.0444 | -0.2183 | -0.0413 |
| b_MutationH526D:SpeciesAbaylyi | -0.1849 | 0.0435 | -0.2688 | -0.1003 |
| b_MutationH526N:SpeciesAbaylyi | -0.1017 | 0.0436 | -0.1854 | -0.0185 |
| b_MutationH526Y:SpeciesAbaylyi | -0.1031 | 0.0452 | -0.1897 | -0.0130 |
| b_MutationI572L:SpeciesAbaylyi | 0.0886 | 0.0458 | -0.0012 | 0.1783 |
| b_MutationI572S:SpeciesAbaylyi | -0.0124 | 0.0464 | -0.1006 | 0.0796 |
| b_MutationS509R:SpeciesAbaylyi | -0.2264 | 0.0484 | -0.3210 | -0.1281 |
| b_MutationS512P:SpeciesAbaylyi | -0.1716 | 0.0464 | -0.2623 | -0.0825 |
| b_MutationV146G:SpeciesAbaylyi | 0.2332 | 0.0448 | 0.1468 | 0.3257 |
| b_SpeciesAbaylyi:Temperature30 | 0.3530 | 0.0226 | 0.3086 | 0.3978 |
| b_MutationD516G:Temperature30 | 0.0868 | 0.0444 | 0.0007 | 0.1755 |
| b_MutationH526D:Temperature30 | 0.3979 | 0.0437 | 0.3139 | 0.4842 |
| b_MutationH526N:Temperature30 | 0.0910 | 0.0446 | 0.0039 | 0.1760 |
| b_MutationH526Y:Temperature30 | 0.1032 | 0.0448 | 0.0141 | 0.1901 |
| b_MutationI572L:Temperature30 | -0.0397 | 0.0452 | -0.1287 | 0.0514 |
| b_MutationI572S:Temperature30 | -0.0086 | 0.0459 | -0.1023 | 0.0793 |
| b_MutationS509R:Temperature30 | 0.2384 | 0.0505 | 0.1389 | 0.3370 |
| b_MutationS512P:Temperature30 | 0.2416 | 0.0466 | 0.1510 | 0.3331 |
| b_MutationV146G:Temperature30 | -0.0273 | 0.0451 | -0.1174 | 0.0614 |
| sigma | 0.0757 | 0.0033 | 0.0694 | 0.0824 |
| Intercept | 0.7269 | 0.0046 | 0.7178 | 0.7363 |
| lprior | -3.1435 | 0.0001 | -3.1438 | -3.1433 |
| lp_ | 332.2377 | 4.4018 | 322.8632 | 339.6354 |

